# Sequence conservation, domain architectures, and phylogenetic distribution of the HD-GYP type c-di-GMP phosphodiesterases

**DOI:** 10.1101/2021.11.05.467447

**Authors:** Michael Y. Galperin, Shan-Ho Chou

## Abstract

The HD-GYP domain, named after two of its conserved sequence motifs, was first described in 1999 as a specialized version of the widespread HD phosphohydrolase domain that had additional highly conserved amino acid residues. Domain associations of HD-GYP indicated its involvement in bacterial signal transduction and distribution patterns of this domain suggested that it could serve as a hydrolase of the bacterial second messenger c-di-GMP, in addition to or instead of the EAL domain. Subsequent studies confirmed the ability of various HD-GYP domains to hydrolyze c-di-GMP to linear pGpG and/or GMP. Certain HD-GYP-containing proteins hydrolyze another second messenger, cGAMP, and some HD-GYP domains participate in regulatory protein-protein interactions. The recently solved structures of HD-GYP domains from four distinct organisms clarified the mechanisms of c-di-GMP binding and metal-assisted hydrolysis. However, the HD-GYP domain is poorly represented in public domain databases, which causes certain confusion about its phylogenic distribution, functions, and domain architectures. Here, we present a refined sequence model for the HD-GYP domain and describe the roles of its most conserved residues in metal and/or substrate binding. We also calculate the numbers of HD-GYPs encoded in various genomes and list the most common domain combinations involving HD-GYP, such as the RpfG (REC–HD-GYP), Bd1817 (DUF3391– HD-GYP), and PmGH (GAF–HD-GYP) protein families. We also provide the descriptions of six HD-GYP–associated domains, including four novel integral membrane sensor domains. This work is expected to stimulate studies of diverse HD-GYP-containing proteins, their N-terminal sensor domains and the signals to which they respond.

**IMPORTANCE:** The HD-GYP domain forms class II of c-di-GMP phosphodiesterases that control the cellular levels of the universal bacterial second messenger c-di-GMP and therefore affect flagellar and/or twitching motility, cell development, biofilm formation, and, often, virulence. Despite more than 20 years of research, HD-GYP domains are insufficiently characterized; they are often confused with ‘classical’ HD domains that are involved in various housekeeping activities and may participate in signaling, hydrolyzing (p)ppGpp and c-di-AMP. This work provides an updated description of the HD-GYP domain, including its sequence conservation, phylogenetic distribution, domain architectures, and the most widespread HD-GYP-containing protein families. This work shows that HD-GYP domains are widespread in many environmental bacteria and are predominant c-di-GMP hydrolases in many lineages, including clostridia and deltaproteobacteria.

## INTRODUCTION

Bis-(3’,5’)-cyclic dimeric guanosine monophosphate (c-di-GMP) was originally discovered as the activator of bacterial cellulose synthase (1) and was subsequently recognized as a universal bacterial second messenger (2-6). In many bacteria, c-di-GMP controls a variety of regulatory processes, including protein and polysaccharide secretion, motility, cell development, and biofilm formation (7-9). In some important bacterial pathogens, c-di-GMP is involved in the regulation of virulence (7-12). The cellular levels of c-di-GMP, which typically range from 40 to 80 nM (13), are tightly controlled by multiple synthases (diguanylate cyclases), containing the GGDEF domain, and c-di-GMP hydrolases (phosphodiesterases) that are found in two structurally distinct forms of the EAL and HD-GYP domains. All these domains are widespread in bacteria and are often encoded in multiple copies in the same genome. These domains can also be found in catalytically inactive forms which often bind c-di-GMP and/or participate in protein-protein interactions. Some c-di-GMP molecules appear to form an intracellular pool of this compound whereas others function locally without joining that pool (13-15). The structure and catalytic mechanisms of c-di-GMP synthesis by the GGDEF domains and its hydrolysis by the EAL domains have been characterized in much detail (7, 16-21).

The second c-di-GMP-hydrolyzing domain, HD-GYP, has been the subject of far fewer studies and very few comprehensive reviews (22-25). This domain, named after two of its conserved sequence motifs, was initially described in 1999 (3) as a specialized version of the widespread HD phosphohydrolase domain, which contains the metal-binding His-Asp active-site motif (26). In addition to the wide variety of phosphohydrolases (phosphatases, phosphodiesterases, nucleotidases, nucleases, nucleotidyltransferases), including the Cas3 nuclease of the CRISPR-Cas system (27, 28), the HD domain superfamily includes certain hydratases, oxygenases, and lyases (29, 30). Compared to the consensus ∼100-aa HD domain (26), the HD-GYP domain was defined as a longer (166-aa) unit with a large number of additional conserved residues including the characteristic GYP sequence motif (3).

Domain architectures of HD-GYP-containing proteins included its combinations with periplasmic ligand-binding sensor domains, the phosphoacceptor (receiver, REC) domain of the two-component signal transduction system, and the GGDEF domain that was previously identified in diguanylate cyclases and c-di-GMP phosphodiesterases (2). These domain associations indicated the involvement of HD-GYP in bacterial signal transduction (3). Based on the numbers of GGDEF, EAL, and HD-GYP domains encoded in various bacterial genomes, the latter two domains were predicted to perform the same function, namely, serve as diguanylate phosphodiesterases, catalyzing the hydrolysis of c-di-GMP molecules synthesized by the GGDEF domain (3). Two years later, in 2001, a partial sequence logo of the HD-GYP domain was published, highlighting the conservation of the predicted metal-binding residues (4).

Subsequent studies of various proteins containing the HD-GYP domain confirmed its ability to hydrolyze c-di-GMP, either introducing a single nick resulting in the production of a linear 5’-phosphoguanylyl-(3’,5’)-guanosine (pGpG) or hydrolyzing it further to two molecules of GMP (31-40). Some HD-GYP domains also hydrolyzed its sibling 3’,3’-cyclic-GMP-AMP (cGAMP) (41-43) but not c-di-AMP or any other cyclic dinucleotide second messengers (41). In the past several years, crystal structures of HD-GYP domains from four different organisms have been solved (42, 44-46), shedding light on the mechanisms of metal-assisted hydrolysis of c-di-GMP and cGAMP. Like the ‘classical’ HD domain and in agreement with the original predictions (3, 26), HD-GYP is an all-α domain that consists of at least 7 α-helices.

Despite this progress, HD-GYP domain has been far less characterized than GGDEF and EAL domains and was even called their “neglected small sibling” (22). One of the reasons for that is that, unlike GGDEF and EAL domains, HD-GYP domain is not encoded in *Escherichia coli, Salmonella enterica, Bacillus subtilis*, and some other model organisms (3, 4), see https://www.ncbi.nlm.nih.gov/Complete_Genomes/c-di-GMP.html. Another reason is the absence of an appropriate sequence model of the HD-GYP domain in most public domain databases. As a result, HD-GYP domains are often confused with the ‘classical’ HD domains, which hampers the analysis of their domain architectures and obscures their specific functions. This work aims at filling that void by providing a detailed analysis of sequence conservation, phylogenetic distribution, domain architectures, and the c-di-GMP binding mechanisms of the HD-GYP domain.

## RESULTS AND DISCUSSION

### Sequence model and structure of the HD-GYP domain

The original description of the HD-GYP domain (3) included a 166-aa alignment that, in addition to the HD motif, included many additional conserved residues including the conserved HHExh(D/N)GxGYP sequence motif, where *h* indicates a hydrophobic acid residue and x – any residue (3). This alignment was subsequently used as the basis for the sequence logo representations of the HD-GYP domain in the reviews of c-di-GMP metabolism [Fig. 1c in reference (4) and Fig. 3c in reference (7)]. Similar alignments appeared in the papers describing the crystal structures of HD-GYP domains from *Bdellovibrio bacteriovorus, Persephonella marina, Pseudomonas aeruginosa*, and *Vibrio cholerae* (42, 44-46). **Figure 1** presents an expanded structure-based sequence alignment that captures the key conserved elements of the HD-GYP domain. This alignment features an unusually high number of conserved residues; functional roles of some of them, deduced from HD-GYP domain structures (42, 44-46), are listed in **Table 1**. This 180-aa alignment is longer than most previous ones and includes three C-terminal α-helices, which, despite their common presence in HD-GYP domains, show little sequence conservation (Figure 1). In multi-domain proteins, the C-terminal HD-GYP domain is often preceded by the S-helix (47); the N-terminal HD-GYP domain of the *Aquifex aeolicus* protein Aq_2027 (GenBank accession number AAC07788.1), which was used in the initial description of the HD-GYP domain (3), is also preceded by an α-helix (not shown in Figure 1).

**Table 1.**
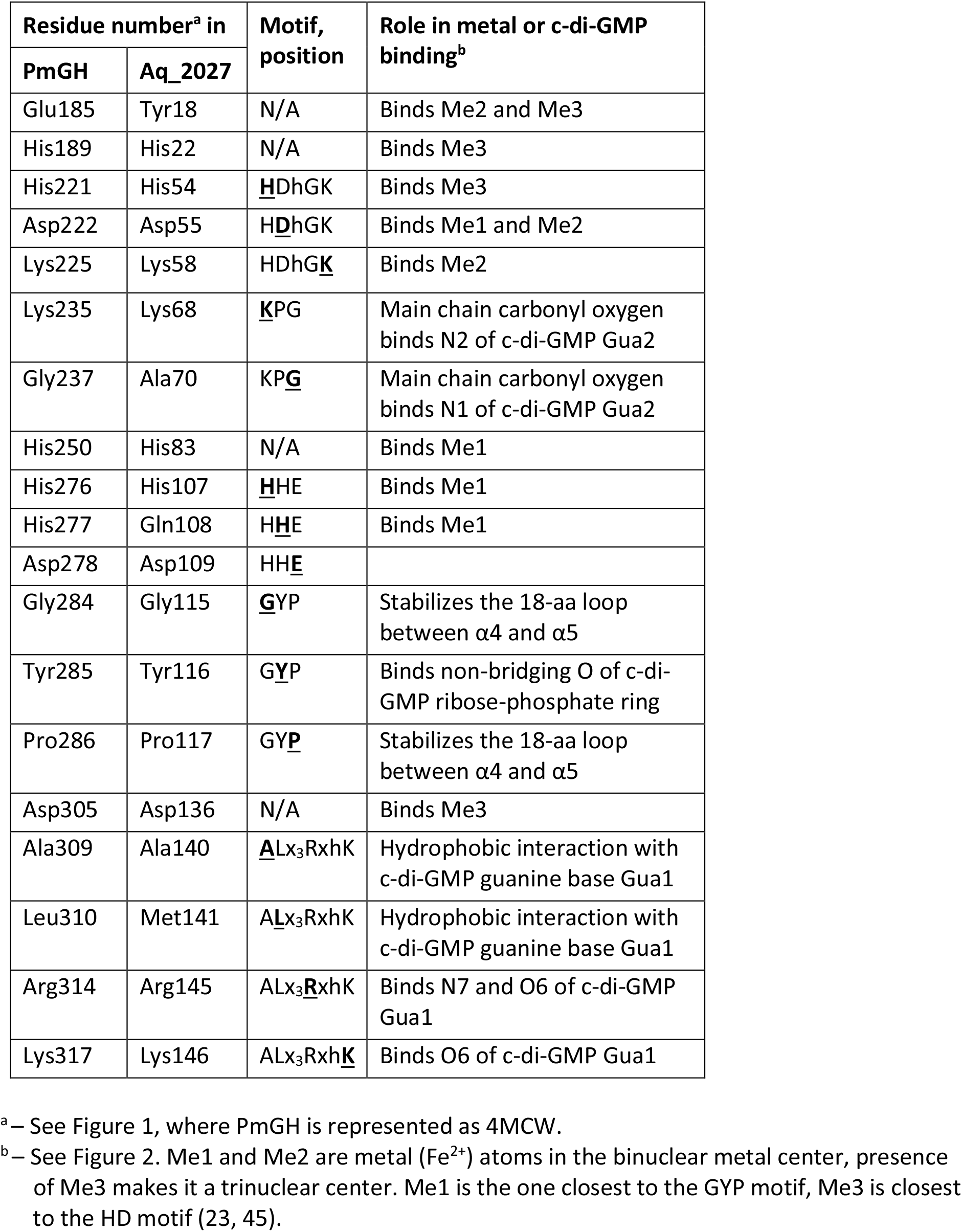
Functions of the conserved residues in the HD-GYP domain.

**Figure 1.**
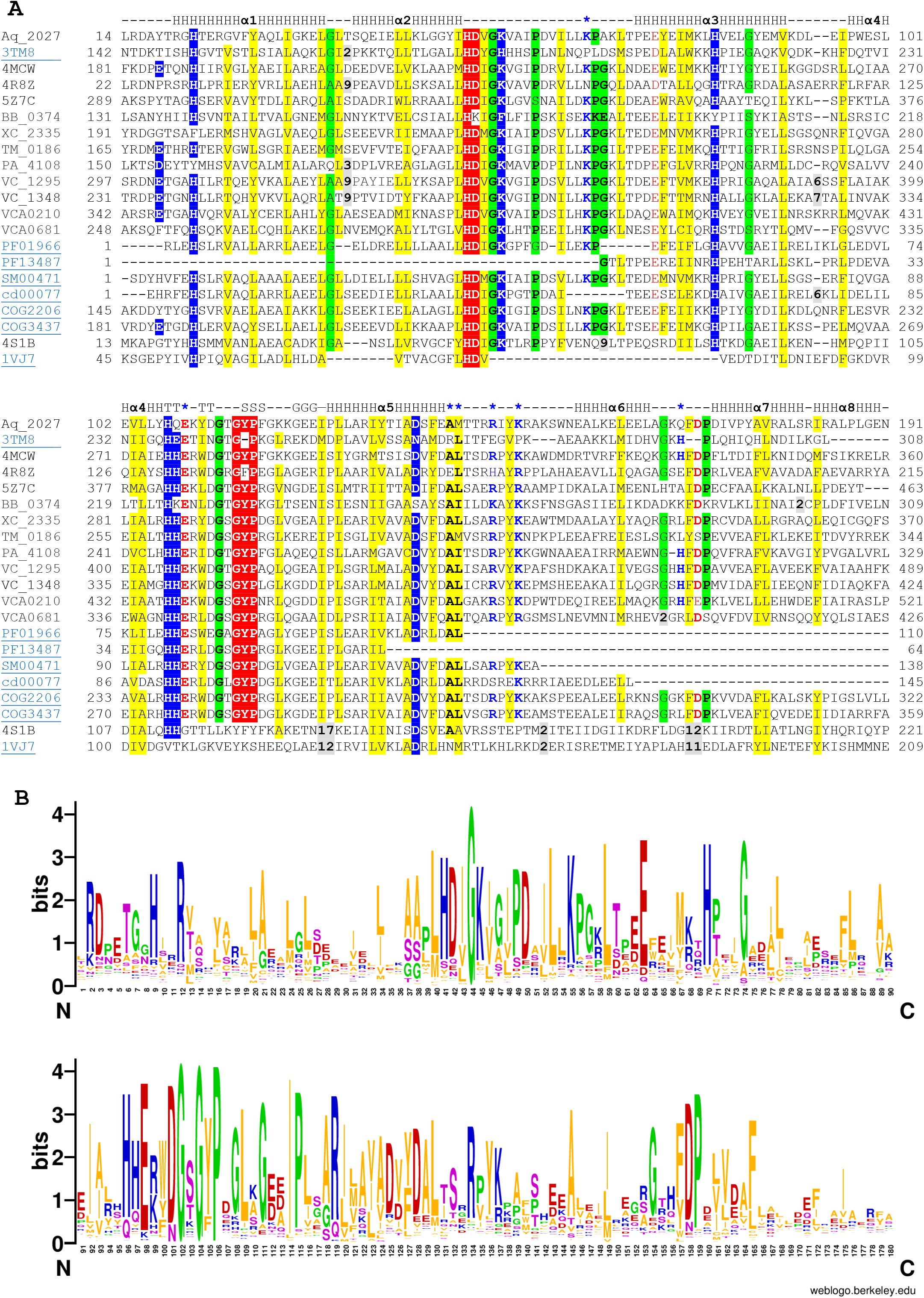
Structure-based sequence alignment and sequence logo of the HD–GYP domain. **A**. The alignment shows the sequences of the *Aquifex aeolicus* protein Aq_2027, used in the initial description of the HD-GYP domain (3), several experimentally characterized HD-GYP domains, and HD-GYP domain entries in several public databases. Amino acids of the eponymous HD and GYP motifs are shown in white on red background. Other metal-binding residues are in white on blue background. C-di-GMP binding residues of PmGH are shown in blue and indicated by asterisks on the top line. Conserved hydrophobic residues are shaded yellow, conserved turn residues (Pro, Gly, Ala, Ser) are shaded green. The first line shows the consensus secondary structure of the domain (H, α-helix; G, 310-helix; T, turn; S, bend). The proteins are listed under their PDB IDs or genomic locus tags; their GenBank accession numbers and respective publications are listed in Table 3. For comparison, two bottom lines show the sequences of structurally characterized ‘classical’ HD (not HD-GYP) domains from two signal transduction proteins: c-di-AMP phosphodiesterase PghP (PDB: 4S1B, CAC99544) (89) and (p)ppGpp synthase/hydrolase RelA (PDB: 1VJ7, CAA51353) (102). **B**. Sequence logo of the HD-GYP domain, derived from an alignment of 1080 sequences of the RpfG (REC–HD-GYP) family.

Unfortunately, most current protein domain databases lack a proper sequence model of the HD-GYP domain. For example, the Pfam database (48) contains two entries for HD-GYP-related domains, HD (PF01966) and HD_5 (PF13487), neither of which covers the entire sequence of HD-GYP. As shown in Figure 1, the 113-aa PF01966 sequence model misses the N-terminal loop that often carries a metal-binding acidic residue (Glu or Asp) and three C-terminal α-helices of HD-GYP. Some sequences in PF01966 contain the GYP motif and likely represent HD-GYP domains, while others do not. Judging from the conservation of the key active-site residues (Figure 1), full-fledged HD-GYP domains comprise at least 8,500 out of nearly 50,000 members of Pfam’s “full” list of HD (PF01966) domains. The 64-aa PF13487 sequence model misses the eponymous HD sequence motif and three other metal-binding residues. Still, an analysis of full-length sequences of ca. 7,700 members of Pfam’s “full” list of HD_5 (PF13487) domains shows that at least 80% of them represent HD-GYP domains. Pfam domain entries HD_2 (PF_12917), HD_3 (PF13023), HD_4 (PF13328), and HD_6 (PF18019) do not seem to include any HD-GYP domains.

Other protein domain databases also have problems in recognizing the HD-GYP domain. The 124-aa HDc sequence model [SM00471](http://smart.embl-heidelberg.de/smart/do_annotation.pl?BLAST=DUMMY&DOMAIN=00471) in the SMART database (49) and its derivatives HDc (cd00077) in the Conserved Domain Database (CDD)(50) and IPR003607 in the InterPro (51) cover the metal-binding residues but lack the C-terminal α-helices. The cd00077 profile also misses the conserved Lys residue (Lys235 in the 4MCW structure, Figure 1) that is involved in c-di-GMP binding. By contrast, the two entries in the COG database (52), COG3437 and COG2206, cover the entire HD-GYP domain but contain additional N-terminal sequences from the receiver (REC, PF00072 in Pfam) domain in the former and the DUF3391 (PF11871) domain in the latter (see below for details). As a result, the HD-GYP entry PS51832 in the ProSite database (53) and its InterPro derivative IPR037522 are the only ones that include the entire HD-GYP domain without any extra amino acid residues. As of October 1^st^, 2021, this InterPro entry IPR037522 contained ∼80,000 entries including 9 crystal structures of the HD-GYP domain from the Protein DataBank (PDB). The structure-based sequence alignment shown in Figure 1 and its expanded version shown in **Figure S1** in the Supplemental file 1 should allow devising suitable sequence models for distinguishing true HD-GYPs from other HD superfamily domains, such as HDc (26) or HDOD (54, 55). As a first step towards this goal, Supplemental file 2 provides a suitable query sequence of the HD-GYP domain and an associated position-specific scoring matrix (PSSM) for use in PSI-BLAST (56) searches and a hidden Markov model (HMM) for use in HMMer (57) searches of protein sequence databases.

### Substrate binding and hydrolysis by HD-GYP domains

Studies of the c-di-GMP- and cGAMP-specific phosphodiesterase activities of the HD-GYP-containing proteins (listed in **Table S1** in the supplemental materials) revealed the key role in catalysis of two metal atoms, usually Fe^2+^ or Mn^2+^, and the metal-binding residues, highlighted in Figure 1 and listed in Table 1 (41, 45). These include the partially conserved Glu residue in the N-terminal loop (E185 in the structure of *P. marina* PmGH, PDB ID: 4MCW, Figure 1). This acidic residue (Glu or Asp) participates in binding the third metal atom; phylogenetic analyses showed that, depending on the presence or absence of this Glu residue, HD-GYP domains could be split into two families carrying, respectively, trinuclear or binuclear metal centers (45, 46, 58). For their c-di-GMP phosphodiesterase activity, experimentally studied HD-GYP domains typically require two ferrous (Fe^2+^) ions, oxidation of either one of them inactivates the enzyme (36). The phosphodiesterase activity has also been observed with manganese (Mn^2+^) ions (36, 39). Some HD-GYP domains could bind Co^2+^, Ni^2+^, Mg^2+^, Zn^2+^, Ca^2+^, and Cu^2+^ with similar affinities (*Kd* around 1 μM), and some of these metal complexes showed comparable catalytic activity (39, 42, 46, 58), reviewed in (25).

The presence of the E185 residue and, accordingly, of the trinuclear metal center, is an important determinant of the ability of the HD-GYP domains to catalyze the second reaction, hydrolysis of pGpG to GMP. In several independent studies, E185-carrying HD-GYP domains of such proteins as VC1295 and VCA0210 from *V. cholerae*, PmGH from *P. marina*, TM0186 from *T. maritima*, PA4108 from *P. aeruginosa*, and DVUA0086 from *Desulfovibrio vulgaris* were shown to hydrolyze c-di-GMP all the way to GMP (36-39, 45), whereas closely related enzymes missing that residue (e.g. containing a Tyr at that position) were unable to do so. However, this requirement is not absolute: PA4781 and VCA0681 proteins, which lack Glu or Asp at that position (Figure 1), have been seen producing GMP as well (35, 37, 38). On the other hand, the isolated HD-GYP domain of the *D. vulgaris* protein DVU0722, which has this Glu residue (Figure S1), was reported to hydrolyze c-di-GMP mostly to pGpG and produce very little GMP, at least in the presence of Mn^2+^ (39). Apparently, there are multiple ways of arranging the HD-GYP domain active site to hydrolyze pGpG and there are additional controlling mechanisms.

As for the substrate, the PmGH–c-di-GMP complex [PDB ID: 4MDZ, (45)] shows a c-di-GMP monomer in a V-shape conformation (59). Its binding involves several conserved amino acid residues that are listed in Table 1 and highlighted in purple in Figure 1. Most of these residues interact with just one of the guanine bases (Gua1 in **Figure 2**), while the second one, Gua2, is primarily bound by main chain carbonyl oxygens of Lys and Gly residues of the highly conserved KPG motif (Lys235 and Gly237 in the PmGH structure, Figures 1 and S1). This organization of the active site accounts for greater flexibility of the Gua2 base and may tolerate its replacement by adenine, which would make the HD-GYP domain able to hydrolyze cGAMP in addition to or instead of c-di-GMP (41-43). The cGAMP hydrolysis could also result from mutations in the Gua1-interacting Arg314-Lys317 pair that forms highly conserved RxxK (or, more precisely, RxY[K/R]) motif (Figure S1) and is largely responsible for the HD-GYP’s specificity for c-di-GMP. In several proteins, including Bd2325, Gmet_3476, and Moth_2495 (Figure S1), replacement of Lys317 with Gln or Asn (resulting, respectively, in the RxxQ or RxxN motifs) shifted substrate specificity towards cGAMP (43). However, HdgA (Vnz_24090, GenBank accession number APE23802.1) protein from *Streptomyces venezuelae* (60), in which the Lys residue of the RxxK motif was replaced by Ser (Figure S1) was still able to hydrolyze c-di-GMP (Dr. Natalia Tschowri, personal communication). In some deltaproteobacterial proteins, this motif is RxxH, with Lys residue replaced by His.

**Figure 2.**
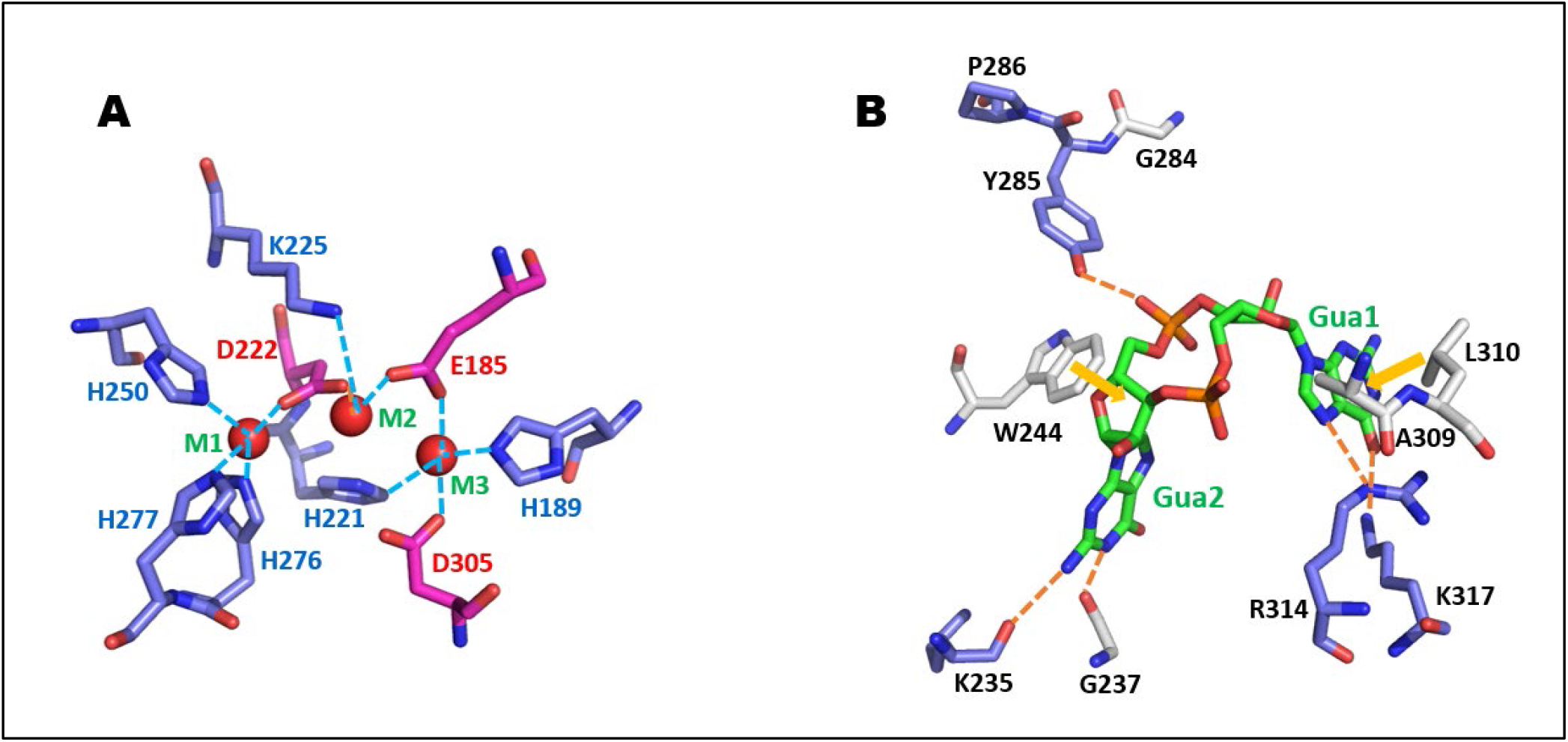
Organization of the active site of the HD-GYP domain. Amino acid residues involved in maintaining the trinuclear metal site (A) and c-di-GMP binding (B) in *Persephonella marina* protein PmGH [PDB ID: 4MCW (A) and 4MDZ (B) (45)]. Residue and guanine base numbering as in Figure 1 and Table 1. Carbon atoms of positively charged residues (His, Arg, Lys) are in blue, those of acidic residues (Asp, Glu) are in pink, and those of uncharged residues are in gray; nitrogen atoms are in dark blue, oxygens are in red. In A, Fe^2+^ ions are shown in dark red, metal coordination bonds are shown as dashed cyan lines. In B, carbon atoms of c-di-GMP are in green, phosphorus atoms are in orange, and oxygens are in red. Hydrogen bonds are shown as dashed orange lines, hydrophobic stacking is indicated by solid yellow lines.

The oxygen atoms of the central 12-member ribose-phosphate ring of c-di-GMP (and cGAMP) are bound by Me^2+^ ions that are coordinated by the metal-binding residues of HD-GYP. The Tyr285 residue of the eponymous GYP motif enhances the binding strength and substrate specificity of the HD-GYP domain by forming a hydrogen bond with one of the non-bridging oxygen atoms of the ribose-phosphate ring (Figure 2).

The catalytic mechanism of c-di-GMP hydrolysis by HD-GYP domains is believed to involve a nucleophilic attack on the phosphorus atom of the central ribose-phosphate ring by a hydroxide ion, activated by one or two Fe^2+^ (or other Me^2+^) ions of the active site (36). The mechanistic aspects of the phosphodiesterase activity of the HD-GYP domain and its allosteric regulation are described in detail in recent reviews (24, 25).

### Phylogenetic distribution and abundance of the HD-GYP domain

The HD-GYP domain is often assumed to be far less widespread than the isofunctional EAL domain (24). However, this impression is largely due to the *E. coli*-centric view of the bacterial world. Indeed, while the HD-GYP domain is not encoded in the genomes of *E. coli* or *B. subtilis* (3, 4), it is abundant in many environmental bacteria. In the current release of the COG database, which aims at covering microbial diversity (52) and includes complete genomes of 1,187 bacterial species from 1,116 genera (typically, a single representative genome per bacterial genus), HD-GYP domains are found in 651 bacterial genomes, compared to 754 that encode EAL domains (**Table S2** in the supplemental materials). Close to a half of bacteria in the list, 586 of 1,187, encode more EAL domains than HD-GYP domains but 252 genomes encode as many or more HD-GYPs than EALs, and 349 do not encode either of them.

Among the organisms covered by COGs, the number of the HD-GYP domains per genome varies from zero to 26 and does not show a clear correlation with the genome size (**Figure 3**). HD-GYP domains show prevalence among certain representatives of the phyla *Chloroflexi, Dictyoglomi, Planctomycetes, Spirochaetes, Synergistetes*, and *Thermotogae*, as well as in *Clostridia* and the proteobacterial classes *Deltaproteobacteria, Oligoflexia*, and *Zetaproteobacteria* (Table S2). **Table 2** lists some bacterial species with a substantial prevalence of HD-GYP domains over their EAL counterparts. Some of the organisms with multiple HD-GYP domains are extremophiles: thermophiles, halophiles, and alkalophiles, which suggests certain robustness of their ability to hydrolyze c-di-GMP. This trend is seen, for example, in two extremely haloalkaliphilic clostridia, *Halanaerobium hydrogeniformans*, isolated from Soap Lake, WA, and *Natranaerobius thermophilus*, isolated from Wadi An Natrun lake in Egypt. The genome of the former encodes 22 HD-GYP domains and just two EAL domains, the latter encodes 15 HD-GYPs and no EALs (Table 2). As noted previously (3), the syphilis-causing spirochete *Treponema pallidum* encodes three HD-GYP domains and no EALs. Genomes of several apparently benign gut symbionts such as *Adlercreutzia equolifaciens, Lawsonia intracellularis, Sutterella megalosphaeroides*, and *Veillonella parvula*, encode single HD-GYP-containing enzymes as their only c-di-GMP-hydrolyzing phosphodiesterases. Also interesting is the presence of HD-GYP domains, but not EAL domains, in the ancestral cyanobacteria-like organism *Candidatus Melainabacteria* bacterium MEL.A1 and the early-branching cyanobacterium *Gloeobacter violaceus* PCC7421.

**Figure 3.**
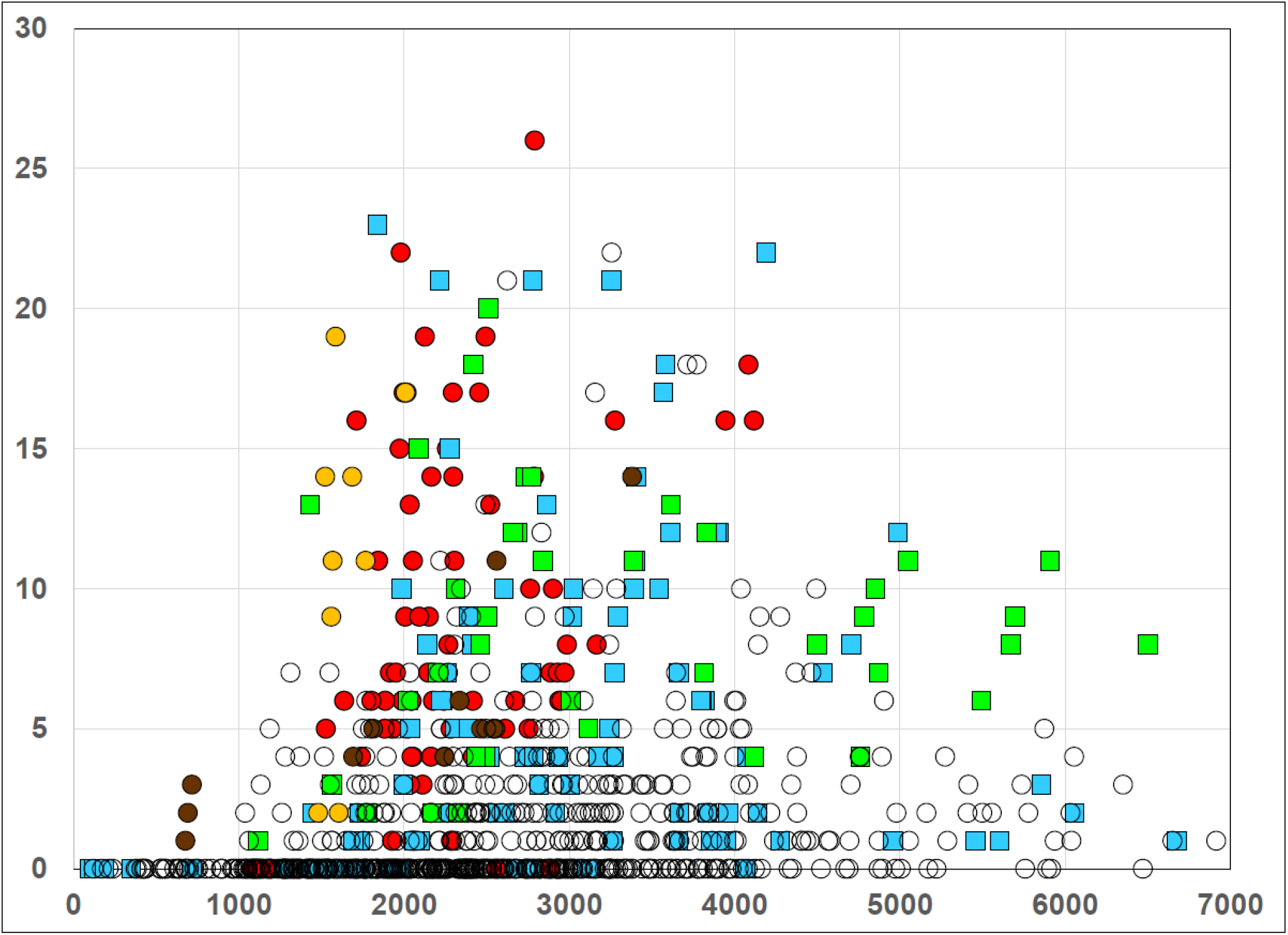
HD-GYP domain content of various bacteria. The numbers of HD-GYP-containing proteins encoded in selected bacterial groups as listed in the COG database (52). The symbols represent members of the following taxa: blue squares, *Betaproteobacteria*; green squares, *Deltaproteobacteria*; red circles, *Clostridia*; yellow circles, *Thermotogae*; dark brown circles, *Spirochaetes*; empty circles, other bacteria.

**Table 2.**
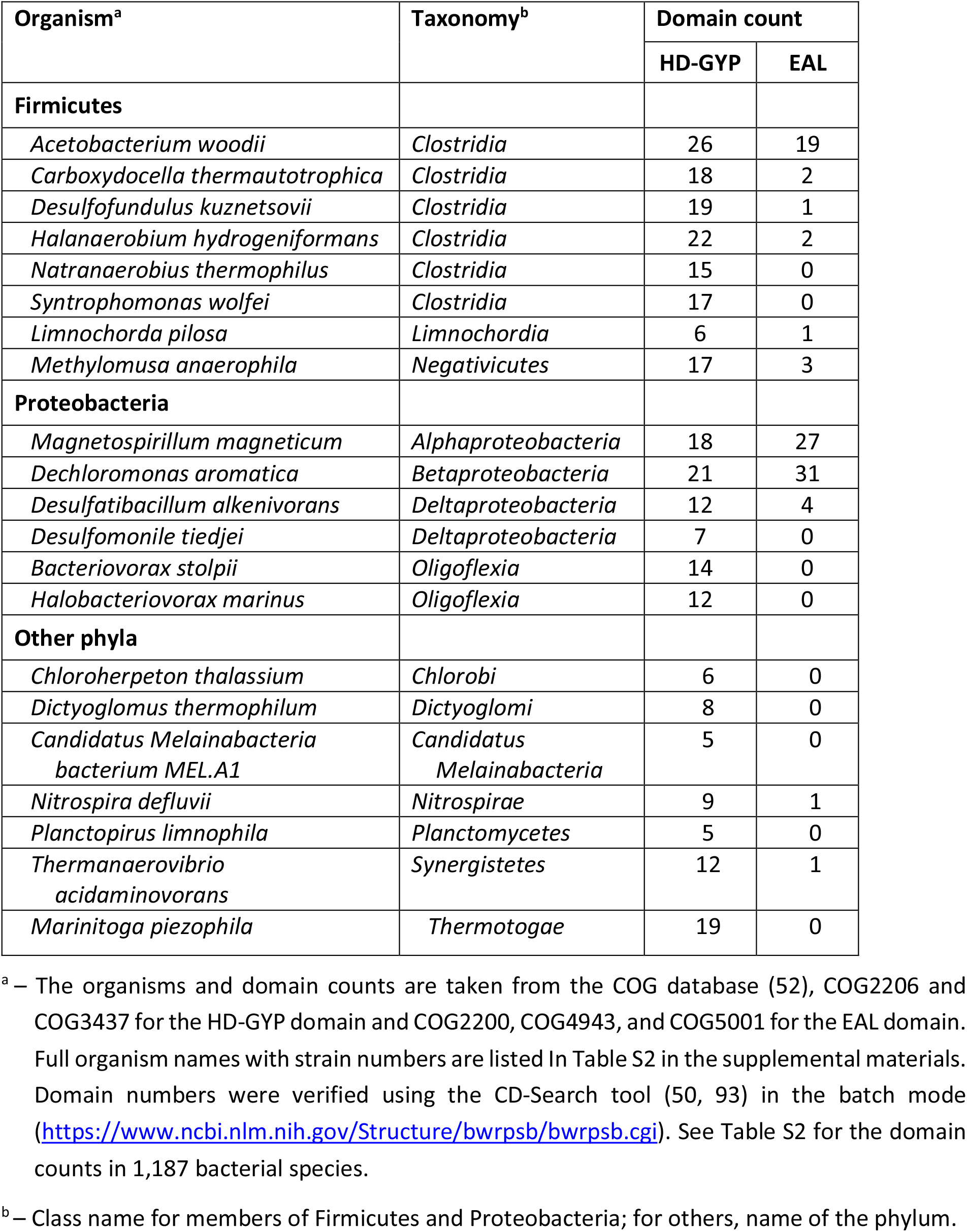
Selected organisms with high numbers of HD-GYP domains.

As c-di-GMP signaling machinery appears limited to bacteria, HD-GYP domains, like GGDEF and EAL domains, were previously reported to be absent in archaea and eukaryotes (7). The only reported exception was a single ORF (RCIX2274, GenBank accession number CAJ37382) from the methanogen *Methanocella arvoryzae* MRE50 that combines REC, PAS, PocR, GGDEF, and HD-GYP domains. The current study found in GenBank 17 eukaryotic (**Table S3**) and 70 archaeal ORFs (**Table S4**) that contained HD-GYP domains. Three HD-GYP-encoding genes from plants had introns, supporting their likely eukaryotic origin. However, one of these ORFs, AXX17_ATUG04560 from *Arabidopsis thaliana* Landsberg *erecta* (61), was absent from the reference sequence of *A. thaliana*. Most eukaryotic HD-GYP-containing proteins were ≥99% identical to certain bacterial proteins (Table S3). Among archaeal HD-GYPs, only the above-mentioned *M. arvoryzae* ORF came from a complete genome, all others were encoded by metagenome-assembled genomes (MAGs) and represent cases of potential bacterial contamination, as discussed, for example, in reference (62). A case in point is a protein with the HD– HD-GYP domain architecture (GenBank accession number HGZ60150) from *Fervidicoccus fontis* MAG (63) that is not encoded in the complete genome of *F. fontis* strain Kam940 (64). Future studies will show if any of the proteins listed in Tables S3 and S4 are genuine archaeal or eukaryotic ORFs.

### Domain architectures of HD-GYP-containing proteins

The most common domain architecture involving the HD-GYP domain is its combination with the two-component receiver (REC) domain, first described in 1999 (3), and subsequently characterized in the nearly identical RpfG proteins from *Xanthomonas axonopodis, Xanthomonas campestris*, and *Xanthomonas oryzae* (31, 65, 66, 67). These proteins were shown to regulate biofilm formation and virulence to citrus, crucifers and rice, respectively. The reported c-di-GMP phosphodiesterase activity was fairly low {Ryan, 2006 #211, 68, 69) (the first of these papers was later retracted), in agreement with the absence of a key His residue in their sequences (Figures 1 and S1). However, a recent report of robust c-di-GMP hydrolysis by a very similar RpfG protein (Figure S1) from *Lysobacter enzymogenes*, another member of *Xandomonadaceae* (70), suggests that this His residue might not be essential after all. The proteins with the REC–HD-GYP domain architecture are referred to as the RpfG family of response regulators (71), see **Table 3**. The experimentally characterized RpfG family members from *Thermotoga maritima, V. cholerae*, and *Desulfovibrio vulgaris* contain the full set of active site residues (Figures 1 and S1) and have robust c-di-GMP phosphodiesterase activity (34, 36, 38, 39). Catalytically inactive proteins of the RpfG family participate in regulation through their protein-protein interactions (see below).

**Table 3.** Domain architectures of selected HD-GYP-containing proteins.

| Family name<sup>a</sup>, domain architecture<sup>b</sup> | Examples (locus tag, GenBank accession number) | References |
| --- | --- | --- |
| RpfG family,<br>REC – HD-GYP | XC2335 (CAB89843)<br>TM0186 (AAD35279)<sup>c</sup><br>VC1348 (AAF94506)<sup>c</sup><br>DVU1181 (AAS95659)<br>DVUA0086 (AAS94372)<br>SO2541 (AAN55571)<br>Gmet_3476 (ABB33684)<sup>d</sup> | (31, 34, 36, 38, 39, 41, 43) |
| Bd1817 family,<br>DUF3391 – HD-GYP | Bd1817 (CAE79676)<br>BB0374 (AAC66766)<br>PA4108 (AAG07495)<sup>c</sup><br>TP0912 (AAC65863)<br>TM1699 (AAC65727)<br>VC2497 (AAF95639)<br>XAC0350 (AAM35242) | (33, 37, 41, 44, 68) |
| PmGH family,<br>GAF – HD-GYP | PERMA_0986 (ACO03551)<br>Tlr0485 (BAC08037)<br>TDE2659 (AAS13176)<br>VC2340 (AAF95483) | (38, 45, 83) |
| VcGAP1 family,<br>HD – HD-GYP | VCA0681 (AAF96581)<sup>cd</sup><br>VCA0931 (AAF96828)<sup>d</sup><br>Moth_2495 (ABC20777)<sup>c</sup><br>SO3491 (AAN56484) | (32, 38, 41-43, 58) |
| RpfX family,<br>REC – PAS – HD-GYP | DVU2933 (AAS97405)<br>SYN_01234 (ABC78345) | (39) |
| RpfY family,<br>REC – DUF3369 – HD-GYP | DVU0722 (AAS95202)<br>VCA0210 (AAF96122)<sup>cd</sup> | (38, 39, 41) |
| PmxA family,<br>dCache – HAMP – HD-GYP | MXAN_2061 (ABF85902)<sup>c</sup><br>TM1682 (AAD36749) | (43, 101) |
| HdgA family,<br>MASE9 – HD-GYP | TM1467 (AAD36535)<br>Vnz_24090 (APE23802) | (60) |
| MASE10 – HAMP – HD-GYP | VC1295 (AAF94454)<sup>c</sup><br>Saut_1462 (ADN09509) | (38, 41) |
| MASE11 – HD-GYP | SO1387 (AAN54452)<br>EW093_05120 (QEN04106) |  |
| MASE12 – HD-GYP | Bas09147 (AAT53240) | (81) |
| HD-GYP – GGDEF | Aq_2027 (AAC07788)<br>BLW93_04085 (OMH40677) |  |
| PAS – HD-GYP | Dgeo_0663 (ABF44965)<br>Glov_3421 (ACD97126) |  |
| PBPb – HD-GYP | Dtur_1688 (ACK42960)<br>TM1170 (AAD36245) |  |
| CHASE2 – HD-GYP | Apau_0390 (EFQ22824)<br>Dpep_1277 (EFC91303) |  |
| Unc – HD-GYP | Bd2325 (CAE80147)<br>TP_0764 (AAC65727)<br>Vnz_24095 (APE23803) | (43, 60) |

<sup>a</sup> – The families are named after their best-characterized member, if available.
<sup>b</sup> – Domain names are from the following entries in the Pfam or SMART databases: REC, PF00072 or SM00448; DUF3391, PF11871, see text and Figure S3; GAF, PF01590; PAS, PF00989; DUF3369, PF11849, see text and Figure S4; dCache, PF02743; HAMP, PF00672; MASE9, MASE10, and MASE11, see Figures S5, S6, and S7; GGDEF, PF00990; PBp, PF00497 or SM00062; CHASE2, PF05226; Unc, uncharacterized domain. Domains with multiple sequence models, such as PAS\_1..PAS\_9 or GAF\_1..GAF\_3 were combined.
<sup>c</sup> – These proteins were able to hydrolyze c-di-GMP to GMP.
<sup>d</sup> – These proteins were able to hydrolyze cGAMP (41, 43).

A fair number of bacterial response regulators have more complex domain architectures with REC and HD-GYP domains separated by one or more PAS or GAF domains. The only experimentally studied protein with the REC–PAS–HD-GYP domain architecture is DVU2933 from *D. vulgaris*. This protein could not hydrolyze c-di-GMP (at least without phosphorylation of the REC domain) but its isolated HD-GYP domain showed Mn^2+^-dependent catalytic activity (39). The *dvu2933* gene is part of an operon that regulates the *lpxC* gene, which is involved in lipid A biosynthesis, but its specific role is unknown (39). In Table 3, proteins with REC–PAS–HD-GYP domain architecture are referred to as response regulators of the RpfX family, in keeping with the RpfG designation for the REC–HD-GYP domain architecture. The central PAS (or GAF) domain is expected to modulate signal transduction from the phosphoacceptor REC domain to the HD-GYP domain, controlling the c-di-GMP phosphodiesterase activity of the protein.

The HD-GYP domain is also involved in a variety of other domain combinations, see Table 3 for selected examples. As noted above, owing to the absence of the standard sequence model for the HD-GYP domain, distinguishing these domains from ‘classical’ HD domains could be challenging. However, analysis of the domain architectures of these proteins revealed a clear distinction: while ‘classical’ HD-domains are typically associated with house-keeping enzymatic activities, HD-GYP domains are usually involved in c-di-GMP-dependent signaling and associated with other signaling domains, either sensory (ligand-binding) or signal transduction domains, such as REC, GAF, PAS, GGDEF, and/or GGDEF-EAL pair. Indeed, a CD-search showed that out of 1,320 response regulators listed in Pfam as combining REC and HD (PF01966) domains and 1,010 sequences with the REC–HD_5 domain combination, at least 90% contain full-length HD-GYP domains, i.e. belong to the RpfG family. With the GAF domain, the picture was the same: all domain combinations listed in Pfam as GAF–HD, GAF_2–HD, GAF_3–HD, GAF_2–GAF–HD, GAF_2–GAF_2–HD, GAF–HD_5, and GAF–HD–HD_5 were found to contain C-terminal HD-GYP domains. Likewise, sequence analysis of the DUF3391 domain (see below) showed that all its combinations with HD and HD_5 domains listed in Pfam represent genuine HD-GYP domains. This was also true for one-component signaling proteins combining the HD domain with periplasmic sensory domains, initially described in (3). In all 29 cases of the PBPb–HD domain combination, 18 out of 19 cases of CHASE2–HD domain combinations, and all 33 cases of dCache_1– HD, dCache_1–HD_5, and dCache_1–GAF–HD_5 domain combinations, listed in Pfam, the output domain proved to be HD-GYP. Associations with PAS domains were not so clear-cut: all PAS_3–HD, PAS_9–HD, and PAS_9–PAS_3–HD entries contained HD-GYP domains, whereas in PAS–HD and PAS_4–HD combinations, up to a quarter of the sequences represented either ‘classical’ HD domain or an inactivated HD-GYP that has lost most of its metal-binding residues.

### Novel HD-GYP-associated domains

A great majority of HD-GYP domain architectures in Pfam (48) show HD or HD_5 as the only recognized domain in the entire protein. The remaining portions of these proteins can be expected to harbor other, highly diverged, poorly characterized, or unrecognized protein domains. Here we provide descriptions of two such domains, which are particularly widespread and are referred to as <u>d</u>omains of <u>u</u>nknown function, DUF3391 and DUF3369, in Pfam (48). We also describe four novel integral membrane domains that are likely to be sensing diverse ligands to regulate c-di-GMP hydrolysis by downstream HD-GYP domains.

### DUF3391

The high-resolution crystal structure of *B. bacteriovorus* Bd1817 (PDB ID: 3TM8 (44)) revealed a 120-aa N-terminal domain, formed by four β-strands with a 310 helix between strands 3 and 4, followed by two long α-helices (**Figure S3**); it is connected to the HD-GYP domain of Bd1817 by a long S-helix. In Pfam, this domain is referred to as Domain of Unknown Function (DUF3391, PF11871). The DUF3391–HD-GYP domain combination, referred to as the Bd1817 family in Table 3, is widespread in bacteria. Among others, it is found in spirochetes and comprises the only HD-GYP-containing c-di-GMP phosphodiesterase in most *Borellia* spp., including the Lyme disease-causing *B. burgdorferi* (33), and one of three such proteins in *Treponema pallidum*, the causative organism of syphilis (72). It should be noted, however, that while the *Treponema* proteins contain full-size DUF3391 domain, the proteins from *Borellia* spp., including the experimentally characterized BB_0374 (33), are missing the first 40 amino acid residues of DUF3391 (Figure S3A). The DUF3391–HD-GYP domain combination has been seen in two other experimentally characterized proteins, PA4108 from *P. aeruginosa* and VC2497 from *V. cholerae*, both of which hydrolyzed c-di-GMP to pGpG and could also convert pGpG to GMP (37, 38). Further, PA4108 showed catalytic activity when expressed in *E. coli* (37), indicating that the DUF3391 domain did not interfere with the activity of the downstream HD-GYP domain and could promote formation of the active, dimeric form of the enzyme. Many genomes contain multiple proteins of the Bd1817 family, thus, *Shewanella* spp. often contain three or even four such proteins (Figure S3A) (Martin-Rodriguez *et al*., submitted). An analysis of the domain architectures of DUF3391-containing proteins showed that almost all of them featured the DUF3391–HD-GYP combination.

The 136-aa HMM logo of the DUF3391 domain that is available on the Pfam website https://pfam.xfam.org/family/PF11871#tabview=tab4 shows only a couple of conserved residues, Gly12 and Trp22. However, this could be attributed to the presence of variable-length Pro-rich inserts between α1 and α2, whose lengths range from zero in Bd1817 to 48 amino acid residues in GNIT_0984 protein from *Glaciecola nitratireducens* (Figure S3, UniProt entry G4QJZ9_GLANF). Collapsing these low-complexity inserts revealed a conserved core structure with several more conserved residues. Unfortunately, the sequence of Bd1817 is so far from the domain consensus (Figure S3B) that it does not allow making any predictions regarding ligand-binding surface(s) of DUF3391. The structural prediction by trRosetta {Yang, 2020 #287} shows the positions of the most conserved residues, including the hydrophobic stacking of Phe29 with Phe35 and potentially also Phe15 and His27 (Figure S3C). However, the roles of other residues, including the highly conserved Met14, remain obscure. It remains to be seen if DUF3391 binds its ligand(s) with the flat surface of its three-stranded β-sheet or through specific interaction with Met14 and other conserved residues.

### DUF3369

(PF11849 in Pfam) is a widespread GAF-like cytoplasmic domain that is frequently associated with the HD-GYP domain and can also be found in diguanylate cyclases, sensor histidine kinases, and hybrid GGDEF–EAL domain proteins (**Figure S4**). Sequence analysis of Pfam entries with DUF3369–HD and DUF3369–HD_5 domain combinations showed that almost all of them contain full-length HD-GYP domains at their C-termini and N-terminal REC domains (sometimes, diverged ones), so they could be considered response regulators of the two-component signal transduction system (54). Two proteins with the REC–DUF3369–HD-GYP domain architecture (the RpfY family in Table 3) have been experimentally characterized. One, VCA0210 from *V. cholerae*, has been mentioned above as a phosphodiesterase capable of hydrolyzing c-di-GMP to GMP (38). The expression of the *vca0210* gene was regulated by quorum sensing and its overexpression resulted in increased motility and decreased biofilm formation (38). In addition to c-di-GMP, VCA0210 was also able to hydrolyze cGAMP to linear dinucleotide (41). Another experimentally studied protein with the same domain architecture, DVU0722 from *D. vulgaris*, did not appear to hydrolyze c-di-GMP, at least without phosphorylation of the REC domain. However, its isolated HD-GYP domain showed Mn^2+^-dependent catalytic activity, producing pGpG and very little GMP (39). While no crystal structure of DUF3369 has been determined so far, a search with HHpred (73) suggests its structural similarity to GAF. Structural prediction by trRosetta (74), available on the Pfam website shows (48) does not show clustering of the conserved residues of DUF3369 (Figure S4B), so the ligand that it binds, if any, remains obscure. Anyway, like the PAS domain in the RpfX family, the DUF3391 domain in RpfY family proteins likely modulates signal transduction from REC to the HD-GYP domain, controlling its phosphodiesterase activity. Remarkably, as seen in Figure S4A, all four DUF3369-containing proteins of *Desulfomicrobium baculatum* belong to the RpfY family (i.e. have the REC–DUF3369–HD-GYP domain architecture). In contrast, five DUF3369-containing proteins of *Azospirillum lipoferum* are far more diverse: two of them have HD-GYP as the output domain, one has the GGDEF-EAL combination, and two more are histidine kinases with distinct domain architectures (Figure S4A).

### MASE9

The original description of the HD-GYP domain (3) identified its fusions with periplasmic sensory domains, subsequently identified as PBPb (TM1170 protein in *T. marítima*) and dCache (TM1682 protein in *T. marítima*) domains. In contrast, in *T. marítima* TM1467 protein, HD-GYP was linked to an N-terminal integral membrane fragment (3) that had not been characterized until now. Sequence analysis of this N-terminal fragment revealed an integral membrane protein domain with 8 predicted transmembrane (TM) segments, two of which were too short to completely traverse the membrane and formed a likely intramembrane hairpin (**Figure S5**). Five positions were occupied by conserved aromatic residues (Phe or Tyr) and 3 positions by hydrophilic (Asn or Ser) residues. By analogy with the previously defined <u>m</u>embrane-<u>a</u>ssociated <u>se</u>nsor domains (MASE1 through MASE8 (75-77), Martin-Rodriguez *et al*., submitted), this domain was named MASE9. In addition to the MASE9–HD-GYP domain combination, MASE9 was also found in several proteins with the MASE9–HD-GYP–GAF–GGDEF domain architecture, as well as in the association with the GGDEF–EAL domain combination (Figure S5), confirming that it represents a promiscuous sensory domain. The poor conservation of the short periplasmic segments, coupled with the presence of several charged residues in the hydrophobic core of the membrane, suggests that ligand binding by MASE9 occurs within the membrane. The only experimentally characterized MASE9-containing protein was HdgA (Vnz_24090) from *Streptomyces venezuelae*, which was poorly expressed and therefore did not appear to have a major role in the c-di-GMP metabolism of this organism (60).

### MASE10

The VC1295 protein of *V. cholerae* was initially characterized as one of those whose expression was repressed by bile acids (78). In its sequence, the HD-GYP domain is preceded by a predicted transmembrane domain and a HAMP domain (38). VC1295 has been shown to bind c-di-GMP (79) and hydrolyze it to both pGpG and GMP (38). The VC1295 entries in UniProt, Pfam, and SMART databases all predicted its N-terminal domain to consist of 6 TM segments but provided no information on its membrane topology. Sequence analysis identified similar 6TM N-terminal regions in three classes of signaling proteins: c-di-GMP phosphodiesterases with HD-GYP domains, adenylate/guanylate cyclases producing cAMP or cGMP, and Ser/Thr protein phosphatases of the PP2C family (**Figure S6**). The diversity of its output domains identified this 6TM region as a novel integral membrane sensor domain, which was named MASE10. The presence of conserved Cys, Ser, and Trp residues in the periplasmic loops of MASE10, as well as conservation of aromatic residues and four charged (Arg, Asp, Lys, and another Asp) residues in the predicted TM segments, suggest highly specific interaction of MASE10 with its ligand(s). In all three classes of signaling proteins, MASE10 domain was followed by the signal-transducing HAMP domain (80)(Figure S6).

### MASE11

Sequence analysis of *Shewanella oneidensis* protein SO1387 revealed another conserved integral membrane domain (**Figure S7**). It contained 7 predicted TM segments but, as in MASE9, two predicted TM segments were too short to cross the membrane and formed a likely intramembrane hairpin. This domain was named MASE11. In addition to its combination with the HD-GYP domain, MASE11 was found in association with PAS, GAF, and GGDEF–EAL domains, as well as in sensor histidine kinases with various domain architectures (Figure S7). So far, no MASE10-containing protein has been experimentally characterized.

### MASE12

A recent study of the roles of c-di-GMP in *Bacillus anthracis* demonstrated the involvement of the HD-GYP-containing protein Bas0914 (GenBank accession number AAT53240.1) in the regulation of bacterial swarming, albeit not in sporulation, germination, or toxin production (81). This protein, referred to as CdgM, combines HD-GYP with a conserved N-terminal integral membrane domain, hereafter MASE12 domain (**Figure S8**). The MASE12-HD-GYP domain combination is encoded in a number of bacilli but has not been seen outside of *Bacillales*. MASE12 was also found in bacillar histidine kinases, including BA5029, previously identified as one of the sporulation kinases of *B. anthracis* (82). Despite the relatively narrow phylogenetic distribution of MASE12, its presence in (predicted) c-di-GMP phosphodiesterases and histidine kinases indicates that it is a novel sensory domain. The presence of conserved aromatic residues in each of the transmembrane segments of MASE12 suggests that its ligand could be a molecule with one or more aromatic rings.

### Regulatory roles of HD-GYP-containing proteins

HD-GYP domains participate in bacterial regulatory cascades in two principal ways: by hydrolyzing c-di-GMP (and/or cGAMP) and by participating in complex protein-protein interactions (24, 25). Like GGDEF and EAL, the other two domains involved in c-di-GMP metabolism, HD-GYPs are rarely seen in a stand-alone form. As shown in Table 3 and can be seen in domain architecture charts of HD and HD_5 domains in Pfam, HD-GYPs are typically found in at the C-termini of multidomain proteins, fused to various N-terminal domains. By analogy with other signal transduction proteins that combine N-terminal sensor domains with the C-terminal output domain(s), such as histidine kinase, adenylate cyclase, GGDEF, EAL, or GGDEF-EAL pair, N-terminal (sensor) domains are expected to modulate the c-di-GMP phosphodiesterase activity of the downstream HD-GYP domains.

In those HD-GYP domains that are linked to extracytoplasmic (dCache, PBPb, CHASE2) ligand-binding sensor domains (Table 3), c-di-GMP hydrolase activity could be directly controlled by the outside signals. To the best of our knowledge, no such protein has been experimentally characterized so far. Fusions of HD-GYP with PAS and GAF domains likely control c-di-GMP levels in response to the intracellular signals, including the levels of AMP, GMP, the redox status of the cell, and other parameters. One such protein, PmGH, has been structurally characterized (45), but the ligands of its N-terminal GAF domain have not been identified. In another protein of the GAF–HD-GYP domain architecture, Tlr0485 from *Thermosynechococcus vulcanus* (GenBank accession number BAC08037.1), the c-di-GMP phosphodiesterase activity of the HD-GYP domain was dependent upon cAMP binding to its GAF domain (83). Another example of such regulation is the protein FV185_09380 (GenBank accession number KXW56972.1) from the betaproteobacterium *Ferrovum* sp., whose N-terminal hemerythrin domain (PF01814 in Pfam) was shown to function as a redox and oxygen sensor that controls the activity of the downstream HD-GYP domain (84). While there is little doubt that the same regulatory mechanisms work in other HD-GYP-containing proteins, they remain to be directly demonstrated.

The widespread RpfG (REC–HD-GYP) family of response regulators (Table 3) includes both catalytically active and inactive forms. Several RpfG family proteins have been shown to hydrolyze c-di-GMP (34, 36, 38, 39) and their phosphodiesterase activity was dependent on the phosphorylated state of their N-terminal REC domains. Thus, along with the response regulators of the WspR (REC–GGDEF), PleD (REC–REC–GGDEF), and PvrR (REC–EAL) families that catalyze respectively, synthesis and hydrolysis of c-di-GMP, RpfG family proteins require their cognate sensor histidine kinases (or acetyl phosphate) for activity and thereby put cellular levels of c-di-GMP under the control of the two-component signal transduction systems. This is yet another way for extracellular signals to control multiple c-di-GMP-dependent functions.

Catalytically inactive proteins of the RpfG family may still control synthesis and secretion of extracellular enzymes and exopolysaccharides through protein-protein interactions. One example is the eponymous RpfG from *X. campestris* (65, 66, 85, 86), which does not appear to have significant c-di-GMP phosphodiesterase activity. Another protein of the RpfG family, *Shewanella oneidensis* HnoD (SO_2541, GenBank: AAN55571.1) with degenerate active motifs (HD⟶SE, HHE⟶QLE, see Figure S1), did not hydrolyze c-di-GMP by itself and inhibited c-di-GMP hydrolysis by the PvrR family response regulator HnoB (REC–EAL domain combination (34)). Another example is protein ECA3548 (GenBank: CAG76446) from *Pectobacterium atrosepticum*, in which an inactivated HD-GYP domain with altered active-site motifs (HD⟶HT, HHE⟶ANE, see Figure S1) is fused to a YcgR-like combination of N-terminal PilZN (aka YcgR, PF07317) and canonical PilZ (PF07238) domains (87). Such a domain combination should be able to bind c-di-GMP but not hydrolyze it, which is consistent with the absence of either c-di-GMP or pGpG hydrolysis by this protein (40). Overexpression of ECA3548 led to increased c-di-GMP levels and affected cell motility (40). There is little doubt that the list of such catalytically inactive HD-GYP domain proteins that are involved in regulatory interactions will continue to grow

## CONCLUDING REMARKS

This study was prompted by the absence of clear data on the distribution of the HD-GYP domains in the public databases. Fortunately, analysis of the domain architectures of HD and HD-GYP domain proteins revealed a clear distinction between them: ‘classical’ HD-domains were typically associated with either house-keeping enzymatic domains, such as RNA polymerase, RNaseY, helicases, and various nucleotidyltransferases, or with defense systems (Cas3 of the CRISPR-Cas system) (27-30). By contrast, HD-GYP domains were always associated with other signaling domains, either sensory (ligand-binding) or signal transduction domains (Table 3). Moreover, certain widespread domain architectures appeared to be limited to the HD-GYP domains. Indeed, CD-search of various proteins combining the HD-type domains with REC, GAF, PBPb, dCache, CHASE2, or DUF3391 domains showed that almost all of them actually contained HD-GYP domains. The only enzymes with the ‘classical’ HD domains with a role in signal transduction were the (p)ppGpp-hydrolyzing enzymes of the RelA/SpoT Homologue (RSH) family and c-di-AMP-hydrolyzing enzymes of the PgpH family (88, 89), see Figure 1. It should be noted, however, that many HD-GYP-containing proteins have undefined N-terminal domains; such proteins comprise the most common domain architectures for both HD (PF01966) and HD_5 (PF13487) entries in Pfam (48). This work delineated three such domains, MASE9, MAS10, and MASE11 (Figures S5, S6, and S7), but there are definitely many more.

Compared to the ‘classical’ HD domains, HD-GYP domains show conservation of several additional residues, including the eponymous GYP motif (Figure 1), whose central Tyr residue binds a nonbridging oxygen atom bond in the central ribose-phosphate ring of c-di-GMP, see Figure 2. All these residues play a role in metal and/or c-di-GMP binding (Table 1). The involvement of multiple residues in c-di-GMP binding is not unique for the HD-GYP domain and has been seen in such c-di-GMP binding proteins as *V. cholerae* MshEN (90) or *Streptomyces venezuelae* RsiG (91, 92). It is important to note partial conservation of the metal-binding Glu residues in the N-terminal loop (E185 in PmGH, Figure 1). Phylogenetic studies of the HD-GYP domains reveal a clear split between those with binuclear (lacking this Glu residue) and trinuclear (with this Glu) metal sites, with the latter group apparently capable of both steps of c-di-GMP hydrolysis, first to pGpG and then to GMP (36, 45, 46, 58). Remarkably, many organisms, including *V. cholerae, P. aeruginosa*, and *D. vulgaris*, appear to encode members of both families (38, 39), see Figure S1. In the emerging pathogen *Shewanella algae*, four out of seven HD-GYP-containing proteins belong to one family and three to another (Martín-Rodríguez. *et al*. submitted). Thus, if pGpG has its own functional role(s), as suggested by some authors (24, 37), some HD-GYP-containing proteins could play a role in hydrolyzing it as well.

Since the HD-GYP logos, presented in Figure 1B and Figure S2, are obtained from sequences that were retrieved from BLAST searches with catalytically active enzymes as queries, they may introduce a certain bias, showing the key amino acid residues as being more conserved than they are. Indeed, our study revealed a substantial number of HD-GYP domains with various substitutions in the active site residues. Such HD-GYP domains are expected to lack catalytic activity but may still retain the ability to bind c-di-GMP and/or participate in various protein-protein interactions. While proteins with GGDEF and EAL domains often participate in local signaling (13-15), there are very few reports of HD-GYP domain proteins acting this way (34, 40, 66). The full extent of such regulation remains to be determined, it is expected to be most pronounced in organisms where HD-GYP is the dominant c-di-GMP phosphodiesterase (Table 2). We hope that the ability to clearly define the HD-GYP domains, whether fully conserved or not, based on the alignment in Figure S1, would allow further insight into the evolution and functional divergence of the HD-GYP-containing proteins.

In summary, HD-GYP domains represent a specialized, highly conserved, widespread, and apparently ancient branch of the vast HD superfamily. Sequence conservation and the narrow functional repertoire of HD-GYP domains can be contrasted with the extreme divergence of the ‘classical’ HD domains that led to the emergence of numerous enzymes with distinct catalytic activities (26, 29, 30), all but two of them unrelated to signaling (88, 89). The HD-GYP and EAL domain counts, presented in Tables 2 and S2, show a wide distribution of the HD-GYP domains and their presence as the prevalent c-di-GMP hydrolases in many diverse bacteria. We hope that an updated description of the HD-GYP domain, its properties and domain architectures, presented in this work, would stimulate further studies of its regulatory roles, including the spectrum of ligands that affect its c-di-GMP phosphodiesterase activity and/or protein-binding properties.

## MATERIALS AND METHODS

### Identification of HD-GYP domain proteins

Putative HD-GYP domains were identified by searching protein sequence databases with PSI-BLAST (56) and HMMER (57) using the *Aquifex aeolicus* protein Aq_2027 (GenBank accession number AAC07788.1) and the *Thermotoga maritima* protein TM0186 (GenBank accession number AAD35279.1) (3) as queries. Domain architectures of the identified proteins were analyzed using CD-search (50, 93) and/or HHpred (73). These domain architectures were further compared with those in Pfam (48), SMART (49), and InterPro (51) databases. To evaluate the presence of HD-GYP domains in various Pfam domain architectures with HD (PF01966) and HD_5 (PF13487) domain entries, the lists of proteins with respective architectures were downloaded from the Pfam website. Their GenBank/EMBL/DDBJ accession numbers were either taken from the respective protein deflines or retrieved using the UniProt’s Retrieve/ID mapping (https://www.uniprot.org/uploadlists/) tool and used in the CD-search (93) in the batch mode https://www.ncbi.nlm.nih.gov/Structure/bwrpsb/bwrpsb.cgi. The resulting hits against [cd00077](https://www.ncbi.nlm.nih.gov/Structure/cdd/cddsrv.cgi?uid=238032), COG2206, COG3437, PF01966, PF13487, and [SM00471](http://smart.embl-heidelberg.de/smart/do_annotation.pl?DOMAIN=HDc) domain models were individually checked for the alignment length and the presence of the HHExh[D/N]GxGYP sequence motif (*h*, hydrophobic residue; *x*, any residue). The list of experimentally characterized HD-GYP-containing proteins was compiled from literature sources; the presence of HD-GYP domain was verified with CD-search.

### Structure-based alignment of HD-GYP domains

Four distinct structures of the HD-GYP domains were retrieved from the Protein DataBank (PDB) and aligned using DALI (94) and VAST (95) servers. These alignments were then compared to that presented in the original description of the HD-GYP domain (3) and further aligned against [cd00077](https://www.ncbi.nlm.nih.gov/Structure/cdd/cddsrv.cgi?uid=238032), COG2206, COG3437, PF01966, PF13487, and [SM00471](http://smart.embl-heidelberg.de/smart/do_annotation.pl?DOMAIN=HDc) domain models (Figure 1A). Experimentally characterized HD-GYP domains and those from phylogenetically diverse bacteria were added to the expanded alignment (Figure S1) based on their CD-search hits.

### Analysis of HD-GYP-containing domain architectures

The lists of proteins with specific domain architectures were taken from Pfam’s “Domain organisation” links for HD and HD_5 domain entries (48) and/or retrieved from CD-search outputs using CDART (96). For the HD-GYP sequence logo, 325 best hits from the PSI-BLAST (56) search of Aq_2027 sequence against the Pfam’s set of proteins with GAF–HD-GYP domain architecture were retrieved, aligned using the “Flat query-anchored with letters for identities” option of BLAST output, trimmed to fit the sequence consensus shown on Figure 1, and submitted to the WebLogo (97) tool. The same procedure has been used to generate sequence logos of the RpfG (REC–HD-GYP) and Bd1817 (DUF3391–HD-GYP) families.

### HD-GYP and EAL domain counts

The numbers of HD-GYP and EAL domains encoded in specific bacterial genomes were taken from the COG database (52), using combined numbers from COG2206 and COG3437 for HD-GYP and combined numbers from COG2200, COG4943, and COG5001 for the EAL domain. The numbers were verified using the CD-search tool in the batch mode and checking the resulting hits for the alignment length and the presence of the HHExh[D/N]GxGYP sequence motif.

### Analysis of DUF and MASE domains

The alignments of two DUF and three MASE domains were constructed from the PSI-BLAST (56) and HMMER (57) outputs and, where appropriate, manually edited to fit the sequence consensus. The secondary structure of the DUF3391 domain was taken from the PDB entry 3TM8 (44); secondary structure of DUF3369 was predicted using HHPred (73) and JPred (98). For MASE domains, transmembrane segments and their membrane topology were predicted with TMHMM2.0 (99) and compared with UniProt (100) and InterPro (51) entries for the selected proteins.

## ACKNOWLEDGEMENTS

This work was supported by the Intramural Research Program of the National Library of Medicine, National Institutes of Health. We thank Prof. Dr. Natalia Tschowri (Hannover) for helpful comments.

